# Enhanced super-resolution microscopy by combined Airyscan and Quantum-Dot-Triexciton Imaging

**DOI:** 10.1101/2020.05.07.082222

**Authors:** Simon Hennig, Dietmar J. Manstein

## Abstract

Super-resolution fluorescence imaging provides critically improved information about the composition, organization and dynamics of sub-cellular structures. Quantum-Dot-Triexciton Imaging (QDTI) has been introduced as an easy-to-use sub-diffraction imaging method that achieves an almost 2-fold improvement in resolution when used with conventional confocal microscopes. Here we report an overall 3-fold increase in lateral and axial resolution compared to standard confocal microscopes by combining QDTI with the Airyscan approach.

## INTRODUCTION

Access to imaging techniques that resolve structures at the molecular level is now widely available^1 2^. However, imaging below the diffraction limit is still associated with some pitfalls and can be difficult to apply to a specific problem. Quantum-Dot-Triexciton Imaging (QDTI)^3 4 5^ is an easy-to-use, high-resolution confocal imaging method based on the generation and detection of a tri-excitonic (TX) state by successive absorption of three photons in quantum dots (QDs). QD655 are cadmium selenide (CdSe) QDs which, in addition to their use in standard fluorescence microscopy applications with detection of their mono-excitonic (MX) emissions, allow the generation and sensitive detection of higher excitonic states. These higher excitonic states can be readily generated using pulsed or continuous wave (CW) lasers in the range between 350 and 488 nm. The unconventional recombination of triple excitons via the p–p recombination channel^6^, produces a characteristic blue-shifted TX emission line at 590 nm, which is readily separated spectrally from the common s–s recombination channel generating the MX emission line at 655 nm. The detection of TX instead of MX emission leads to increases in lateral and axial resolution adding to the attractiveness of QD655 as a probe for fluorescence imaging.

When QDTI was used in combination with the Airyscan technique^7 8^, we achieved an up to 3-fold improvement in resolution compared to standard confocal imaging. The combined approach, which we refer to as AiryQDTI imaging, allows imaging down to a lateral resolution of 81 nm. The method is easy to use and, apart from Airy filtering, does not require any image post-processing. A single confocal scan is required to generate the super-resolved image. Since AiryQDTI is based solely on the physical effect of triple exciton generation, it is compatible with conventional buffers and requires no elaborate or special sample preparation.

## RESULTS

To determine the resolving power of AiryQDTI imaging, we prepared surfaces that are sparsely decorated with QD655 quantum dots. As the full width at half maximum (FWHM) detected from emitters depends strongly on the applied excitation intensity^3^, we determined the changes in apparent FWHM for individual quantum dots as a function of the excitation intensity. Fig. 1a shows a representative region of interest that was sequentially imaged using Confocal MX, Airyscan MX and Airyscan TX detection. The resulting point spread functions and the dependence of the lateral FWHM on excitation intensity illustrate the up to 3-fold improvement in lateral resolution achieved by applying the AiryQDTI approach (Fig. 1b, c and d). The limitation of the achieved resolution in TX imaging strongly correlates with the excitation intensity in combination with the pixel dwell-time and consequently the signal-to-noise ratio (SNR). When comparing the probabilities of generating and detecting TX emissions via the p–p recombination channel and generating and detecting a MX state in QDs, the probability is much lower in the TX case. This can be compensated for by a higher pixel dwell-time, but has its limitations. Therefore, when the excitation intensity is reduced, the TX SNR decreases faster than the MX SNR, limiting the detection of TX signals in our experiments at about 2.5% excitation intensity instead of 0.2% in the MX case. The respective limits in our experiments are marked by red lines (Fig. 1d).

**Figure 1.**
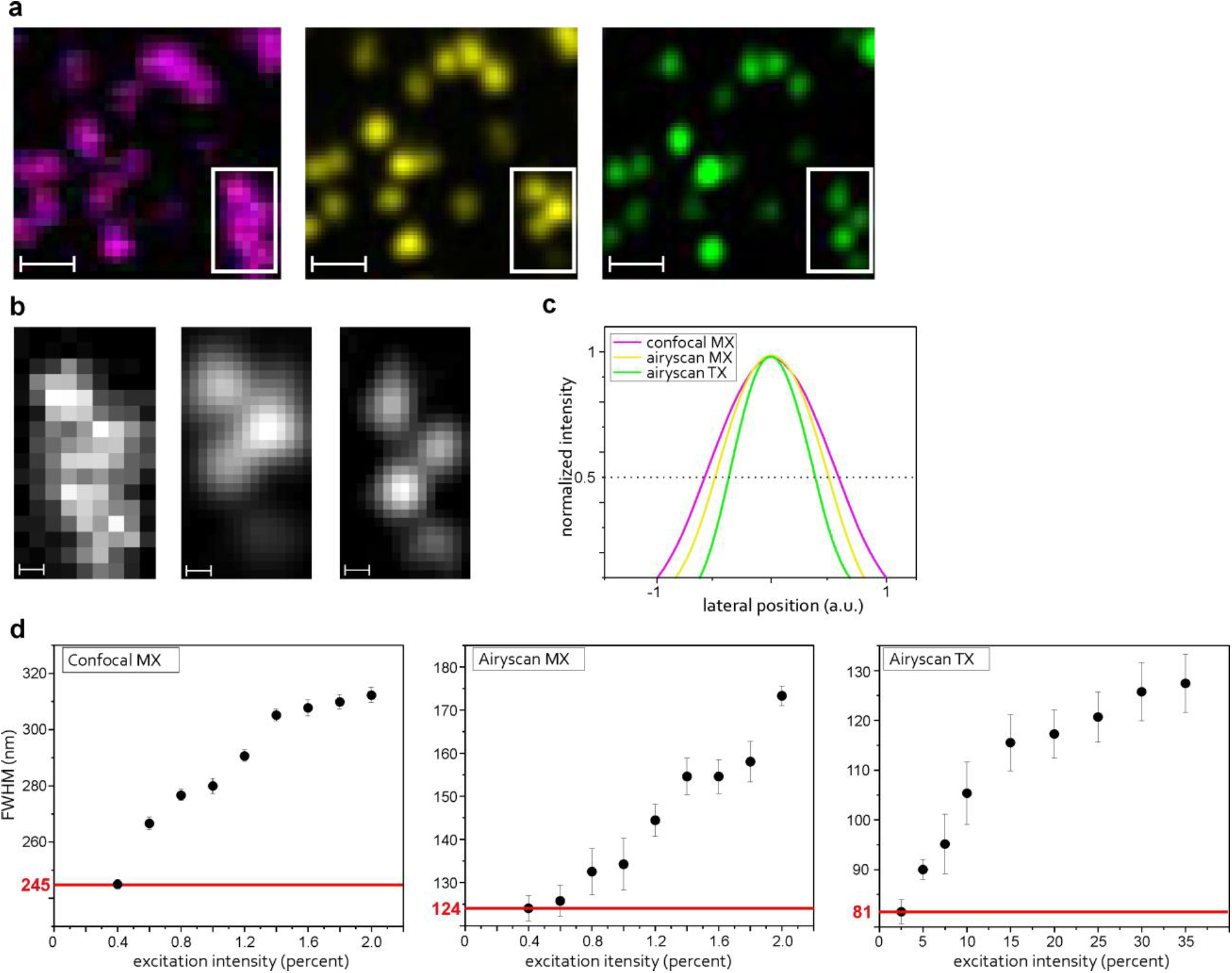
Characterization of AiryQDTI imaging using sparsely decorated QD655 surfaces. (a) The same region of interest showing a surface area sparsely decorated with QD655 quantum dots was imaged using the Confocal MX (magenta), the Airyscan MX (yellow) and Airyscan TX (green) channel. Scale bar, 500 nm. (b) Expanded areas from a (rectangles) showing a cluster of quantum dots in the Confocal MX (left), Airyscan MX (middle), and Airyscan TX (right) channel. Scale bar, 100 nm. (c) Point spread function of a single quantum dot, showing the intensity distribution in the three emission channels. (d) Plots showing the lateral FWHM obtained from a single quantum dot as function of the excitation intensity. All three emission channels show a decreasing FWHM with decreasing excitation intensity. The Confocal MX channel shows a limit at 245 nm, the Airyscan MX channel at 124 nm and the Airyscan TX channel at 81 nm, showing a 2–fold lateral resolution enhancement from Confocal MX to Airyscan MX and 3–fold lateral resolution enhancement from Confocal MX to Airyscan TX. Error bars represent the standard error.

Since the QDTI method improves the resolution in all three dimensions, the AiryQDTI approach leads also to a reduction of the axial FWHM. 3D xz visualization of a sparsely decorated QD655 surface indicates the axial resolution improvement for the Airyscan TX channel, compared to the Confocal MX and Airyscan MX channels (Supplementary Fig. 1). Our estimate of an axial resolution of about 210 nm achieved by AiryQDTI is based on the reported axial resolution of 350 nm for z-stacks recorded in Airyscan super-resolution mode and a further 1.7-fold gain by QDTI^3^. The improved axial resolution results simultaneously in a reduced image background. This is most obvious in images with crowded and densely organized structures, as shown in Supplementary Fig. 2.

**Figure 2.**
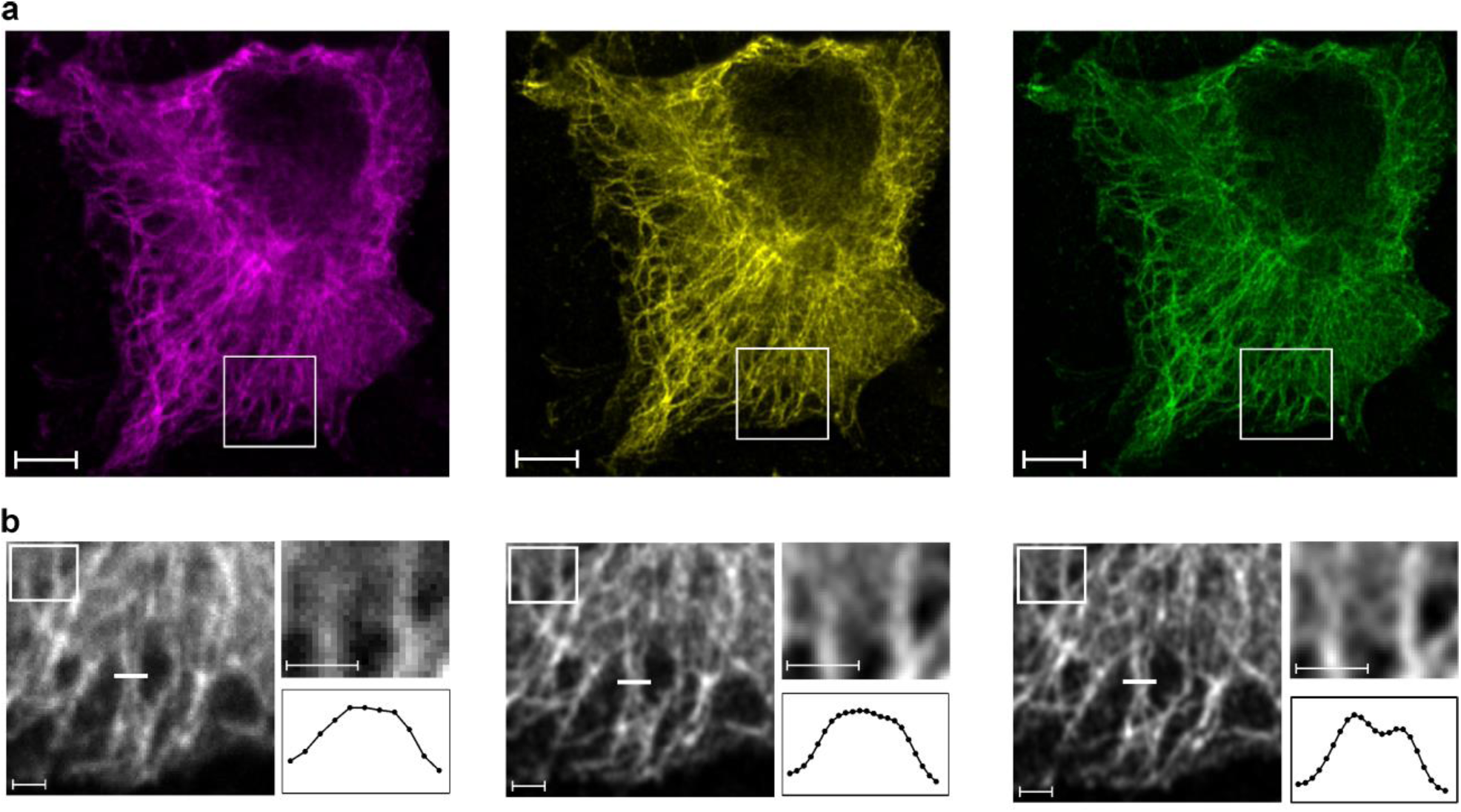
AiryQDTI–mediated improvements in the resolution of cytoskeletal structures. (a) Region of interest showing a fixed A549 cell, immunolabeled for microtubules with QD655 quantum dots and imaged subsequently by utilizing the Confocal MX (magenta), Airyscan MX (yellow) and Airyscan TX (green) emission channel. Scale bar, 5 µm. (b) Insets, taken from a showing the detailed distribution of filamentous structures in the three channels. Fine details can be observed in the Airyscan TX channel, which are not observable in the Confocal MX or Airyscan MX channel (insets). Nearby structures are resolved with greater detail (white lines and associated intensity–profiles). Scale bars, 1 µm.

We show the applicability of the AiryQDTI imaging technique by immunolabeling of microtubule structures in fixed cultured A549 cells (Fig. 2). By scanning with the previously determined optimal imaging conditions for each detection channel, we subsequently generated images from the same region of interest, starting with channels that require low excitation intensities such as Confocal MX and Airyscan MX. The resulting images demonstrate the improved resolution of the Airyscan TX channel compared to the other channels (Fig. 2b, insets and intensity profiles).

## CONCLUSION

In summary, we describe a straightforward and readily applicable confocal imaging technique based on the generation of three exciton states in quantum dots in combination with the Airyscan approach, which can resolve fluorescence signals with a precision of 81 nm laterally and around 210 nm axially. Labeling the structures of interest with QD655 quantum dots in combination with a single confocal scan using moderate illumination intensity and detection of the blue-shifted TX emission with the Airy detector is sufficient to produce the super-resolved image.

## METHODS

Experiments were performed using a Zeiss LSM980 with Airyscan 2 and 34 channel QUASAR detection unit. The principle of the Airyscan detector can be found elsewhere^7^. The 405 nm diode laser of the microscope was employed for excitation of QDs. Confocal MX imaging was carried out with the standard setting of 70 nm/pixel, a detection-window of 630–700 nm, amplification gain 650 V and 2-fold sampling. The pinhole was set to 1 AU. Airyscan imaging was carried out with 30 nm/pixel using 605–705 (MX) and 525–585 nm (TX) filters. Pixel dwell-times were set to 37.66–68.23 µs for Airyscan MX imaging and 78.2–135.3 µs for Airyscan TX imaging with 2-fold sampling. Detector gain was set to 750 V and 950 V, respectively. 3D imaging was carried out with 210 nm/plane for confocal and 120 nm/plane for Airyscan imaging. To avoid image artifacts and to ensure a consistent correction of the images in post-processing with ZenBlue, we used a constant value of 5 for the Airy filter and deactivated the automatic settings. Interpolation was switched off during all image processing steps. Laser intensities were set to 0.4% for MX imaging and 4% for TX imaging in cell images. FWHM measurements were performed in PBS buffer, which contained 1 mM 2-mercaptoethanol to suppress QD655 blinking^9^. Comparison of MX and TX emission was performed by increasing the excitation intensity in 0.2% steps from 0.4% to 2% for MX detection and in 2.5 or 5% steps from 2.5% to 35% for TX detection. Images were processed and analyzed using FIJI^10^. Normalization of images from the three different channels was achieved by using “Contrast Enhancer” (Process>Enhance Contrast) with “normalization”. LUTs and greyscales are linear representations of the raw or normalized values. A549 cells were cultured in DMEM/F-12 medium (Merck KGaA, Darmstadt, Germany). Cells were seeded in LabTek II chamber slides (NUNC) 48 hours prior measurements. The protocol for fixation and immunolabeling of microtubules can be found elsewhere^3^.

## Author Contributions

S.H. designed the project and performed imaging and image analysis. S.H. and D.J.M. interpreted results and wrote the manuscript.

## ACKNOWLEDGMENT

We thank Dr. Rudolf Bauerfeind for his advice and discussions and the MHH Core Unit for Laser Microscopy for support.

