## Supplemental Information for "Enhanced super-resolution microscopy by combined Airyscan and Quantum-Dot-Triexciton Imaging"

### SUPPORTING INFORMATION

SUPPLEMENTARY FIGURES:

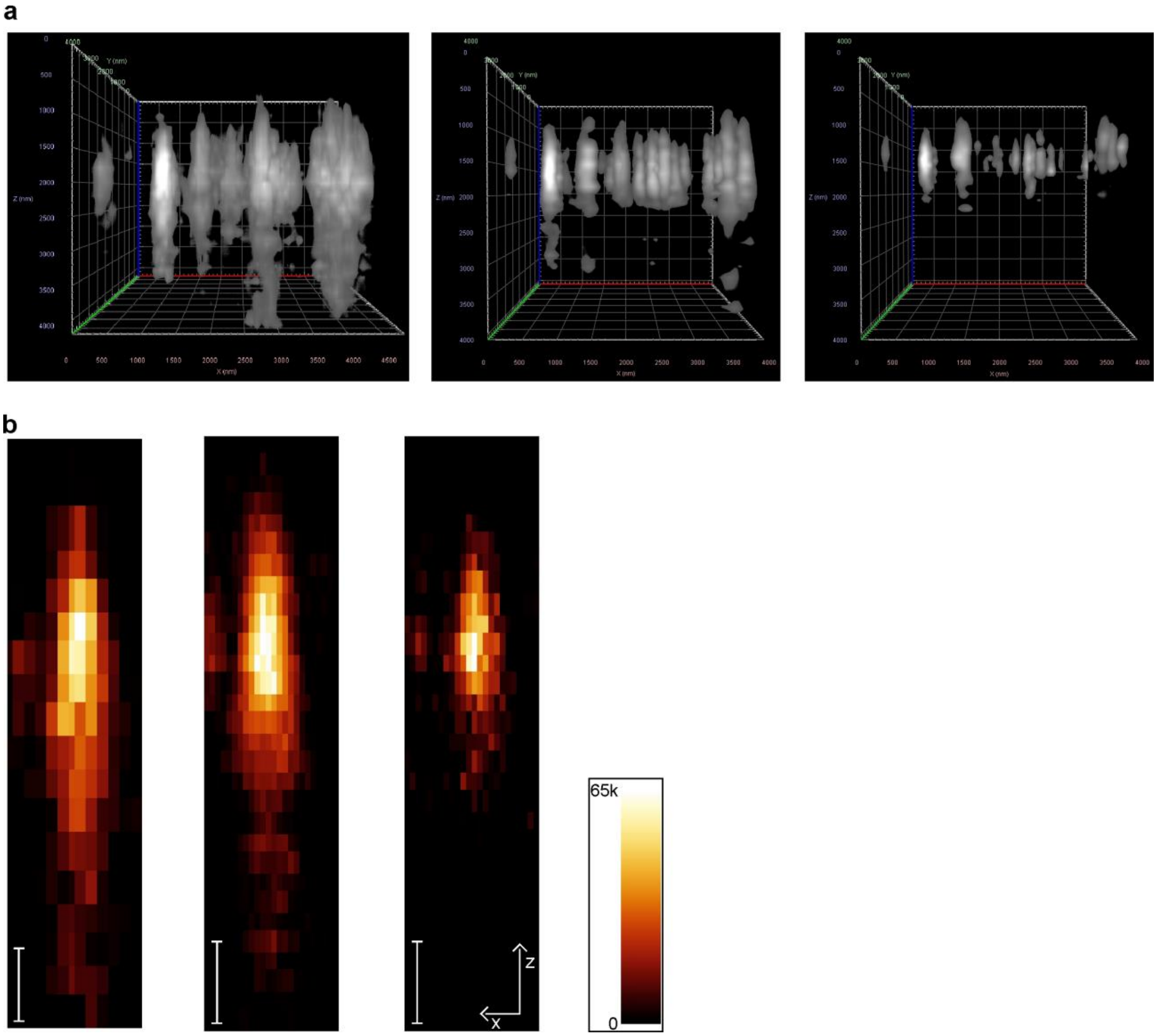


Figure S1. Investigation of Axial Point Spread Functions of QD655 emitters. (a) 3D representation of a glass surface sparsely decorated with QD655 emitters. Channels from left to right: Confocal MX, Airscan MX, and Airyscan TX. Images were generated, using the 3D image view of the ZenBlue software. Image intensities were matched using the min/max function and a setting of 13% for the high-pass intensity filter. (b) Maximum intensity projections of a single QD655 emitter recorded from left to right in the Confocal MX, Airyscan MX and Airyscan TX emission channels. Intensities were normalized as described in the Methods section. Scale bar, 500 nm.

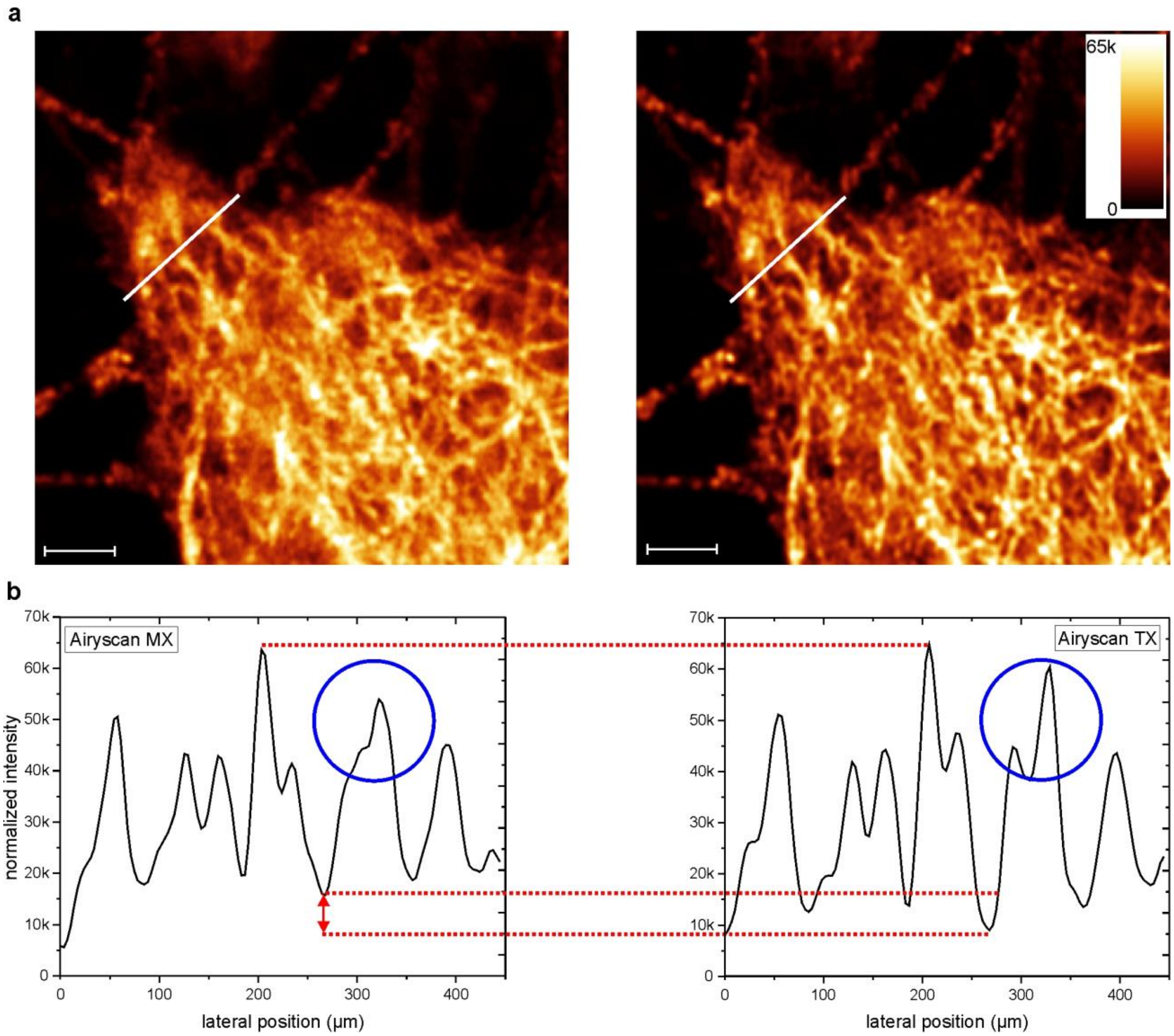


Figure S2. Fluorescence background reduction induced by AiryQDTI. (a) Region of interest showing a dense network of QD655-labeled microtubules imaged in the Airyscan MX channel (left) and Airyscan TX channel (right). Scale bar, 2 µm. (b) Normalized intensity profiles, illustrating the extent to which the background intensity in the Airyscan TX channel is reduced (double headed arrow). The enhanced lateral resolution of the Airyscan TX channel resolves finer details (blue circle).
